# An IDH-independent mechanism of DNA hypermethylation upon VHL inactivation in cancer

**DOI:** 10.1101/2020.12.09.418616

**Authors:** Artem V. Artemov, Svetlana Zhenilo, Daria Kaplun, Alexey Starshin, Alexey Sokolov, Alexander M. Mazur, Justyna Szpotan, Maciej Gawronski, Martyna Modrzejewska, Daniel Gackowski, Egor B. Prokhortchouk

## Abstract

Hypermethylation of tumor suppressors and other aberrations of DNA methylation in tumors play a significant role in cancer progression. DNA methylation can be affected by various environmental conditions including hypoxia. The response to hypoxia is mainly achieved through activation of the transcription program associated with HIF1a transcription factor. VHL inactivation by genetic or epigenetic events, which also induces aberrant activation of HIF1a, is the most common driver event for renal cancer. With whole-genome bisulfite sequencing and LC-MS, we demonstrated that VHL inactivation induced global genome hypermethylation in human kidney cancer cells under normoxic conditions. This effect was reverted by exogenous expression of wild-type VHL. We show that global genome hypermethylation in VHL mutants can be explained by transcriptional changes in MDH and L2HGDH genes that cause the accumulation of 2-hydroxyglutarate—a metabolite that inhibits DNA demethylation by Tet enzymes. Unlike the known cases of DNA hypermethylation in cancer, 2-hydroxyglutarate was accumulated in IDH wild type cells.

**Key points:**

- Inactivation of VHL gene leads to genome hypermethylation in kidney cancer cells. The DNA methylation phenotype can be rescued by endogenous expression of wild-type VHL.
- DNA hypermethylation can be attributed to the accumulation of a Tet inhibitor 2-hydroxyglutarate
- The accumulation of 2-hydroxyglutarate in IDH wild-type cells is explained by gene expression changes in key metabolic enzymes (malate dehydrogenase MDH and 2-hydroxyglutrarate dehydrogenase L2HGDH).

## Introduction

VHL inactivation is the most common driver event for renal cancer [1]. VHL is involved in hypoxia sensing, directing polyubiquitination and further degradation of Hif1a transcription factor [2]. Inactivation of VHL leads to accumulation of *Hif1a* and activation of the genes involved in hypoxia response [3,4]. Therefore, mutation-driven VHL inactivation in tumors causes similar transcriptomic changes as those induced by hypoxic environment [3,5,6]. DNA methylation is sensitive to oxidation-reduction potential. Hypoxia was shown to impair Tet-driven DNA demethylation and decrease the level of hydroxymethyl-cytosine, the key intermediate of the demethylation process, which led to overall DNA hypermethylation [7].

Genome hypermethylation during hypoxia had a wide variety of physiological and molecular consequences. Changes in DNA methylation contribute to pathologies caused by chronic intermittent hypoxia in people living at the sea coast and potentially mediate adaptations to chronic sustained hypoxia by affecting the hypoxia-inducible factor (HIF) signaling pathway [8]. High-altitude acclimatization is thought to have a genetic component, and other factors, such as epigenetic gene regulation, are involved in acclimatization to high-altitude hypoxia in nonacclimatized individuals. Those included significantly lower DNA methylation at high altitudes in EPO and RXRA genes and LINE-1 repetitive elements, and increased methylation in EPAS1 and PPARa genes [9]. In conjunction with these results, it was reported that hypoxia alters the DNA methylation profile of cardiac fibroblasts via Hif1a regulation of DNA methyltransferase DNMT1 and DNMT3B [10]. Apart from its role in environmental adaptations, hypoxia is a major factor that affects epigenetic marks through oncogenic progression. For instance, Hypoxia in LUAD tumor cells led to changes in DNA methylation patterns. FAM20C,

MYLIP and COL7A1 were identified as the hypoxia-related key genes in LUAD progression, which were regulated by DNA methylation [11]. In breast cancers, Mediated by members of the HIF family of transcription factors, hypoxia leads to a more aggressive tumor phenotype by transactivation of several genes as well as reprogramming of pre-mRNA splicing. The expression level of one of the exons of DHX32 and BICD2 significantly correlated with the methylation levels and even made it possible to predict patient survival [12].

It wass therefore obvious that hypoxia could influence genome methylation level. Several non-mutually exclusive mechanisms can be suggested to explain this phenomenon: (i) transcriptional or functional inhibition o TET enzymes directly by oxygen intermediates [7]; (ii) transcriptional of functional enhancement of DNA methylating DNMT enzymes; (iii) de novo activated transcriptional programs that are associated with stabilized nuclear HIF1alpha and cause changes in concentrations of TET/DNMT regulating metabolites. In this work we have explored the possibility of regulating genome methylation level by initiating hypoxia transcriptional programs without changing the actual level of oxygen. Previously, we explored the effects in gene promoters and CpG-rich regions, by performing reduced representation bisulfite sequencing (RRBS) in cells under normoxia conditions but with stabilised HIF1a [13]. However, it remained unknown if the observed changes could be generalized to the whole genome and what mechanism could drive the hypermethylation.

In this work, we profiled genome-wide DNA methylation with bisulfite sequencing (wgBS) in three clones with independent VHL inactivation and observed global genome hypermethylation. Reintroduction of exogenous wild-type VHL has partially rescued DNA methylation phenotype to wild-type state. Global changes in DNA methylation were confirmed with LC-MS. Levels of several metabolites were measured with LC-MS to explore a potential mechanism of global DNA hypermethylation upon VHL inactivation in normoxia.

## Results

### Introduction of cancerous mutation into the VHL gene results in stabilization of HIF1α in Caki1 cells

With CRISPR-Cas9 gene editing, we obtained three clones (Caki1*VHLmut*) of Caki1 human clear cell renal carcinoma cell line with a homozygous frameshift in VHL gene (Figure 1A). The frameshift at 181D led to production of an inactive form of VHL that was unable to ubiqitinate Hif1a [3,13]. Mutations within 181D were frequently observed in renal tumors according to the COSMIC database [14]. Western blot analysis of nuclear extracts confirmed stabilization of Hif1a protein in the absence of hypoxia in cells with mutant VHL in comparison with Caki1 cells (Supplementary figure S1A). Expression of VHL in the three clones (Caki1*VHLexo*) was rescued with exogenous chimeric VHL (tagged with HA) by lentivirus infection. Western blot showed restored VHL expression that led to the degradation of Hif1a protein (Supplementary figure S1A). RNA-seq analysis of Caki1 cell line, three clones with mutant VHL and the clones with rescued VHL expression confirmed the activation of hypoxia program in VHL mutants and its deactivation after exogenous expression of VHL (Supplementary figure S1B).

**Figure 1.**
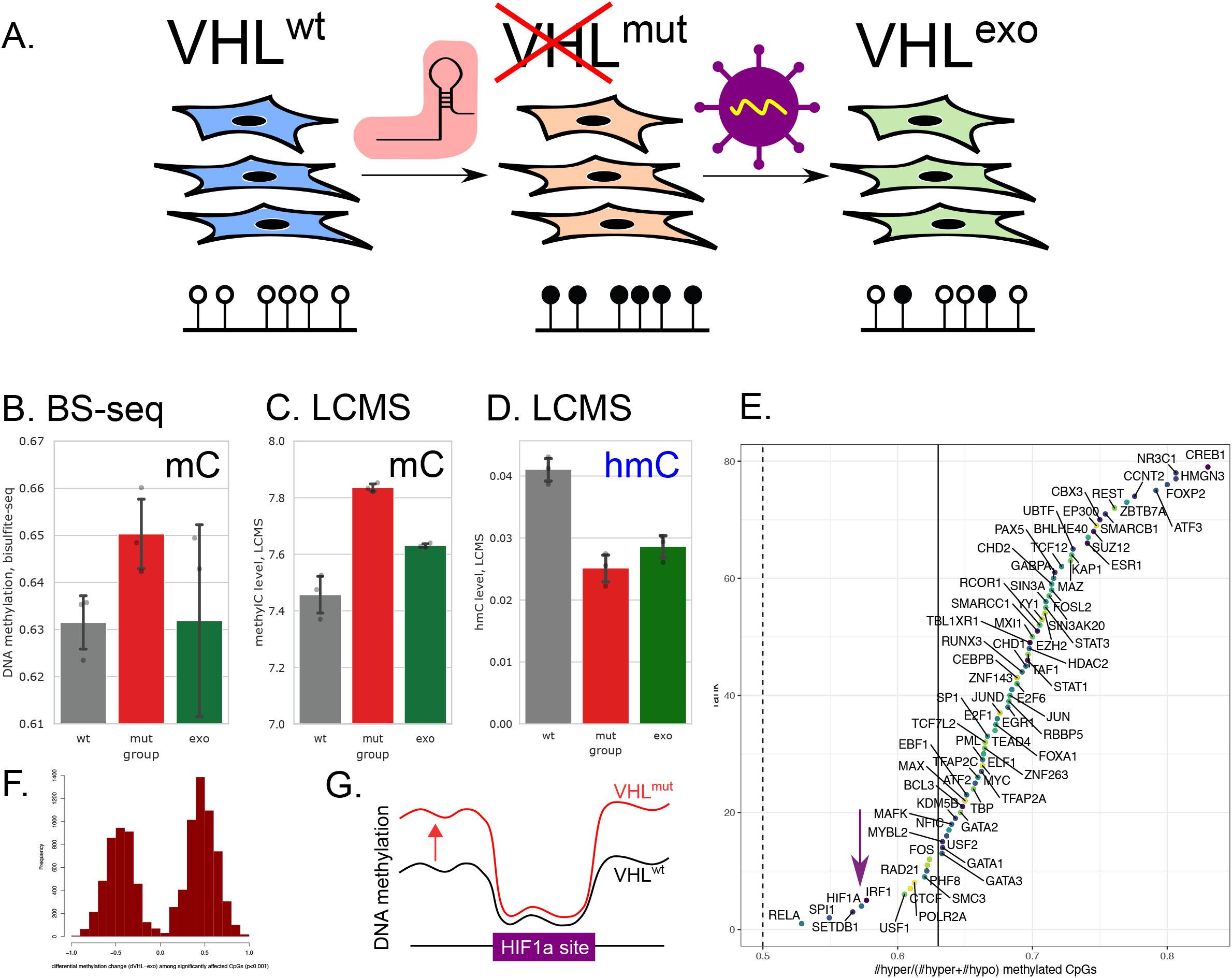
Whole-genome methylation in Caki-1 cells, three clones with inactivated VHL and the same clones with VHL expression rescued by exogenous construct. **A**. Design of the experiment. VHL was inactivated independently in three clones of Caki-1 human kidney cancer cell line. In each clone, the mutation was rescued by exogenous expression of VHL. DNA methylation and metabolites were profiled in these cells. **B**. Global DNA methylation levels estimated from bisulfite sequencing data. **C**. Global DNA methylation levels measured by mass-spectrometry. **D**. Global DNA hydroxymethylation (hmC) measured with mass-spectrometry. **E**. For each transcription factor, we estimated the fraction of hypermethylated sites among all CpG sites with significantly altered methylation. **F**. DNA methylation changes in CpGs with significant difference between wild-type Caki-1 cells and the clones with inactivated VHL. **G**. While all transcription factors tend to be hypermethylated in VHL mutants, HIF1a sites to a great extent avoid hypermethylation.

### VHL inactivation causes whole-genome hypermethylation

To test if the induction of the transcriptional programs driven by HIF1α could cause changes in genome methylation in the absence of hypoxia, we performed whole genome bisulfite sequencing in Caki1, Caki1*VHLmut* and in Caki1*VHLexo* cells (Figure 1A, Supplementary figure S2). In our previous work, we observed DNA hypermethylation in gene promoters. Here we discovered a significant increase of DNA methylation not only in promoter regions, but rather globally throughout the genome (Figure 1B). Hypermethylation could be observed in each chromosome (Supplementary figure S3). The magnitude of the change was smaller for whole-genome DNA methylation (2% increase) than for CpG islands (6.45% increase), as most of the CpG’s in the genome are fully methylated even in the original cell line, hence, DNA methylation levels of these positions could not increase more. Exogenic expression of VHL rescued DNA methylation phenotype and returned global DNA methylation close to the wild-type levels (Figure 1B, S3).

We validated the increase of global DNA methylation in VHL mutants and its partial rescue to the normal phenotype in the clones with exogenic VHL expression by mass-spectrometry (LC-MS) of DNA nucleotides and observed the same effect (Figure 2B, P_mut_vs_wt_ = 0.01). With LC-MS, we were able to distinguish between methylated and hydroxymethylated cytosine in the studied samples. While mC levels increased in VHL mutants, hmC levels decreased (Figure 1D). In Caki1*VHLexo* we saw partial restore in mC which was accompanied by the drop in hmC level. Accumulation of mC accompanied with depletion of hmC could be explained by impaired oxidation of mC to hmC by Tet enzymes — the first step of DNA demethylation [15].

**Figure 2.**
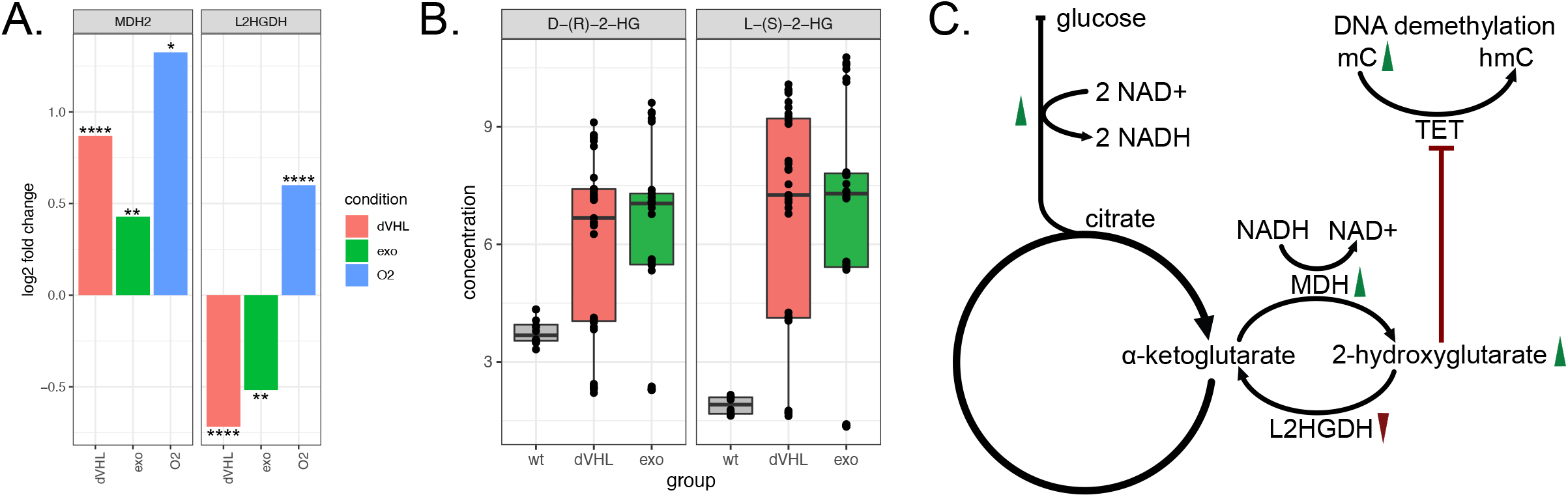
**A**. Changes of gene expressions of MDH and L2HGDH enzymes responsible for 2-hydroxyglutarate synthesis and degradation. Y-axis indicates log fold-change of gene expression in VHL mutants, rescued VHL mutants with exogenic expression of VHL and Caki-1 cells in hypoxic conditions, all compared to Caki-1 cells in normoxic conditions. DESeq2 p-values corrected for multiple testing (FDR) are indicated as follows: **** *P <* 0.0001; *** *P <* 0.001; ** *P <* 0.01; * *P <* 0.05. **B**. Concentrations of L and D isomers of 2-hydroxyglutarate measured by mass-spectrometry. **C**. Proposed mechanism of DNA hypermethylation in VHL mutants. Green arrows indicate the increase of gene expression of all enzymes involved in glycolysis and MDH after VHL inactivation. Red arrow indicates decreased expression of L2HGDH. Increased concentrations of L- and D-hydroxyglutarate in VHL mutants lead to the inhibition of Tet enzymes and DNA hypermethylation.

The observed changes in DNA methylation could not be attributed to the differences in cell cycle progression. Quantitation of DNA content revealed that the differences in the distribution of cells according to the phases of the cell cycle between Caki-1 cells, VHL mutants and the rescued clones were beyond statistical significance (Supplementary Figure S4)

DNA hypermethylation in VHL mutants was not restricted to the binding sites of specific transcription factors or to the loci characterized by certain histone modifications. Among CpG sites with significantly altered DNA methylation in VHL mutants (P_BetaBinomial_<0.01, Figure 1F) that fall into a given set of genomic intervals such as HIF1a binding sites, we checked what proportion of CpG’s were hyper- and hypomethylated. Binding sites of all transcription factors tended to be enriched by hypermethylated CpGs (Figure 1E). Interestingly, one of the smallest rates of hypermethylated CpGs (yet still higher than 0.5) was observed for HIF1a sites. Hif1a binding sites could potentially be protected from hypermethylation by increased binding of Hif1a (Figure 1G).

### Gene expression changes of key metabolic enzymes

To decipher a potential mechanism of the observed DNA hypermethylation in the absence of active VHL, we first studied gene expression of the key enzymes responsible for acquisition of methylation and demethylation. Tet enzymes were transcriptionally downregulated in VHL mutants (Supplementary Figure S5). We checked the protein levels of Tet2 enzymes. Tet2 level was not affected by VHL mutation (Supplementary figure S6A). However, the ability of Tet enzymes to demethylate DNA measured by activity assay kit were not impaired, moreover, some clones demonstrated higher rate of demethylation of the probe DNA (Supplementary figure S6B).

VHL inactivation led to a significant increase in gene expression of the enzymes involved in all stages of glycolysis (Supplementary figure S7). We observed a 3.8-fold decrease of SLC2A1 expression (FDR=2.8*10^-12), 2.2-fold decrease of HK1 (FDR=4.7*10^-4), 2.2 and 2.8-fold decrease of expression for PFKL and PFKP respectively (FDR=1.5*10^-6 and 1.0*10^-19), 3.7- and 7.8-fold decrease for ALDOA and ALDOC (FDR=1.7*10^-21 and 2.3*10^-21 respectively), 3.0-fold decrease for GAPDH (FDR=1.4*10^-13), 4.3-fold decrease for PGK1 (FDR=4.1*10^-21), 2.7-fold decrease for PGAM1 (FDR=2.1*10^-7), 3- and 7.2-fold decrease for ENO1 и ENO2 (FDR=1.9*10^-20 и 3.0*10^-9 respectively) and 3.9-fold decrease for LDHA (FDR=3.9*10^-25).

Glycolysis leads to accumulation of pyruvate and NADH that are subsequently utilized in TCA cycle. Some TCA cycle metabolites are also known to stimulate or inhibit the activity of TET enzymes. With mass-spectrometry, we measured the concentrations of 2-ketoglutarate, fumarate and succinate. No significant changes were observed for 2-ketoglutarate (P=0.87), fumarate (P=0.087) and succinate (P=0.066).

### Accumulation of 2-HG in cells with inactivated VHL

2-hydroxyglutarate can inhibit TET enzymes that perform active demethylation of DNA. In gliomas and acute myeloid leukemias (AML) with mutated IDH1 and IDH2 genes, genome hypermethylation is explained by increased concentration of D-2-hydroxyglutarate that, in turn, was caused by a gain-of-function mutation in IDH [16–18]. However, the studied Caki-1 cells and their derivatives had wild-type IDH1 and IDH2. We validated that IDH1 and IDH2 did not have gain-of-function mutations at positions chr2:209113192 (IDH1), chr15:90631934 and chr15:90631838 (IDH2, coordinates according to hg19) that could lead to aberrant production of D-2-hydroxyglutarate: all positions were covered by RNA-seq data and all reads contained only the wild-type alleles in these positions. We therefore explored what alternative mechanisms could lead to 2-hydroxyglutarate accumulation.

Apart from oxidizing malate to oxaloacetate, malate dehydrogenase (MDH2) is capable of reducing 2-oxoglutarate (TCA cycle metabolite) to L-2-hydroxyglutarate [19,20]. We showed that expression of MDH2 significantly increased (1.8-fold, FDR=2*10^-8) after VHL inactivation compared to the original cell lines with wild type VHL. It significantly decreased after exogenous VHL reactivation (1.3 fold decrease, FDR=0.001, Figure 2A).

In contrast, VHL inactivation caused a two-fold drop of expression of two 2-hydroxyglutrarate dehydrogenases, L2HGDH (0.60, FDR=2.9*10^-5, Figure 2A) and D2HGDH (0.73, FDR=0.03). These enzymes can convert, respectively, L- and D-2-hydroxyglutarate back into 2-oxoglutarate (Figure 2C). Taken together, the observed gene expression changes suggested the increase of 2-hydroxyglutarate and specifically its L-isomer in VHL mutants.

To validate this hypothesis, we measured the isomers of 2-hydroxyglutarate with mass-spectrometry (LC-MS). Indeed, the concentrations of both L- and D-2-hydroxyglutarate significantly increased in VHL mutants and decreased after the rescue of VHL expression (Figure 2B). The L-isomer showed stronger increase (4-fold, P_Wilcoxon_=9.9*10^−5^), whereas the

D-isomer was increased twofold (P_Wilcoxon_=0.0036), which suggested IDH-independent mechanism of 2-hydroxyglutarate accumulation.

To investigate if this increase of 2-hydroxyglutarate concentration was sufficient to explain global DNA hypermethylation in VHL mutants, we profiled 2-hydroxyglutarate levels in HCT116 cell line. D-2-hydroxyglutarate concentration reached 5-7.5 mmol/cell in HCT116 cell line which was within the range of concentrations observed in Caki-1 VHL mutants (Supplementary figure S8). In HCT116 cells, this accumulation of D-2-hydroxyglutarate was sufficient to reach higher levels of DNA methylation than those of Caki-1 VHL mutants.

### Tumors formed by Caki, CakiVHLmut and CakiVHLexo cells

We assessed the ability of the studied clones to form tumors in immunodeficient mice (Figure 3A). All three clones with mutated VHL (Caki1*VHLmut*) gave rise to significantly bigger tumors (more than 2.5-fold increase in tumor size, P_Wilcoxon_=0.001) as compared to the original Caki-1 cells with wild-type VHL. Exogenous expression of wild-type VHL significantly rescued the phenotype in each clone (P_Wilcoxon_=0.0008). In two of the three rescued clones, tumor sizes were not higher than those of Caki1 cells.

**Figure 3.**
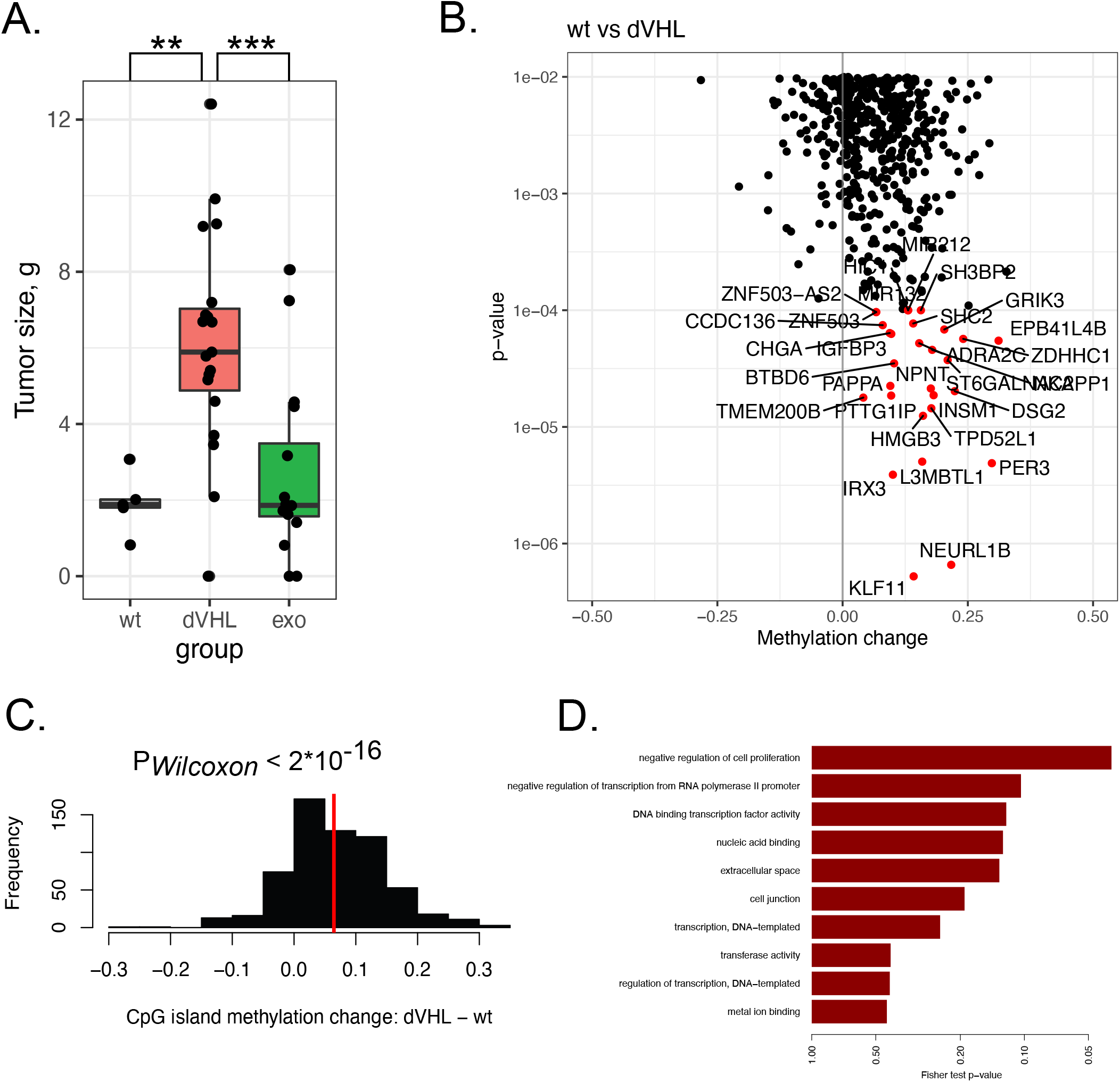
**A**. Mass of the tumors formed by cells injected into immunodeficient mice. VHL mutant Caki-1 cells formed significantly bigger tumors. **B, C**. DNA methylation in CpG islands increases in VHL mutants compared to VHL-wild-type Caki-1 cells. Each promoter CpG island was assigned to a gene. CpG islands were significantly hypermethylated in VHL mutants. **D**. Genes with significantly hypermethylated CpG islands in their promoters were enriched among negative regulators of cell proliferation.

To understand the mechanism of increased tumorigenesis in VHL mutants (Caki1*VHLmut*), we explored DNA methylation in gene promoters containing CpG islands. For each CpG island, we tested if DNA methylation level changed between the original Caki1 cells and VHL mutant cells (Figure 3B). Strikingly, a vast majority of CpG islands were hypermethylated in VHL mutants: on average, a 6.45% increase in DNA methylation level was observed (P_Wilcoxon<2.2*10^-16, Figure 3C). Genes with hypermethylated CpG islands in their promoter regions were strongly enriched among the genes responsible for negative regulation of proliferation (Figure 3D). Among the genes hypermethylated in VHL mutants, we observed known tumor suppressors in kidney cancer: PER3, PCDH19, RLIM, KIAA1024. Their expression was increased in VHL mutants and reversibly silenced upon VHL inactivation (Supplementary figure S9).

## Discussion

DNA hypermethylation in hypoxia was previously linked directly to decreased oxygen levels: active DNA demethylation involves oxidation of mC to hmC by Tet enzymes and therefore is sensitive to oxidation-reduction potential in the cells [7]. However, DNA hypermethylation could be observed at completely normal oxygen concentrations in the cells with impaired hypoxia signalling. We have previously shown that VHL inactivation led to DNA hypermethylation in CpG-rich genomic regions profiled by RRBS [13]. Here, we generalized this observation to the whole genome. We also demonstrate that the rescue of VHL expression from exogenous lentiviral vector partially restores DNA methylation.

Global increase of DNA methylation in VHL mutants and decrease after reactivation of VHL were observed by two independent methods, bisulfite sequencing and mass spectrometry (LC-MS). The magnitude of DNA hypermethylation within CpG islands was even stronger than in the whole genome. Hypermethylation of CpG islands in the promoters of tumor suppressors could explain higher size of tumors formed by cell clones with mutated VHL. Decrease in tumor suppressor gene expression PER3, PCDH19, RLIM, KIAA1024 in VHL mutant cells follows the changes in their promoter methylation. All of these genes are downregulated in various cancer types,regulated by promoter hypermethylation [21–23]. Increased tumor size in VHL mutant cells is also in accordance with increased viability of these cells after achieving confluence in cell culture [3].

Hypermethylated CpG sites were not enriched within specific transcription factor binding sites. However, HIF1a sites were among the least affected by global DNA hypermethylation. We hypothesized that overall genomic hypermethylation is to a great extent compensated within HIF1a sites by increased binding of Hif1a caused by accumulation of Hif1a protein in VHL mutants. Recent findings suggest that HIFs fail to bind if the CpG dinucleotides within their binding sites are methylated, both *in vitro* and *in vivo* [24]. Therefore, the observed protection of the functional Hif1a sites from hypermethylation could prevent their long-term inactivation.

We identified a metabolic mechanism that could lead to DNA hypermethylation in VHL mutants. VHL mutants accumulated 2-hydroxyglutarate, a metabolite that is known to inhibit Tet-mediated DNA demethylation in IDH-mutant glioblastomas. Specifically, the D-isomer of 2-hydroxyglutarate is accumulated in IDH-mutant tumors. However, in the studied cell clones with intact IDH1/2 and mutated VHL, we observed a stronger increase of the concentration of the L-isomer. The D-isomer might arise from tautomerization of the L-isomer and both isomers can subsequently inhibit Tet [25–27]. In addition, we observed an increased expression of MDH enzyme that produces L-2-hydroxyglutarate and decreased expression L2HGDH that degrades it. These observations are in line with L-2-hydroxyglutarate production by MDH, rather than D-2-hydroxyglutarate production by IDH1 or IDH2. Thus, for the first time we demonstrate an IDH-independent mechanism of 2-hydroxyglutarate accumulation that leads to increased DNA methylation upon VHL inactivation. Further studies are required to show how Hif1a can directly or indirectly regulate the expression of MDH and L2HGDH.

Typically, heterozygous mutations within the active sites of isocitrate dehydrogenases (R132H in IDH1 and R140Q in IDH2) lead to the accumulation of millimolar concentrations of R-2-hydroxyglutarate. However, even the tumors with wild type IDH1 and IDH2 can accumulate higher levels of 2HG (0.01-0.1 mM) than normal tissues (10^−8^ M) [28]. To demonstrate that the magnitude of the increase of 2-hydroxyglutarate concentration was sufficient for the observed DNA hypermethylation, we measured the concentrations of L- and D-2-hydroxyglutarate and the levels of DNA methylation in HCT116 cell line. In HCT116 cell line, the similar or lower concentrations of L- and D-2-hydroxyglutarate induced even higher levels of DNA methylation compared to the studied VHL mutants.

In previous works, a difference in the concentration of 2-hydroxyglutarate was observed between various cell lines and tumors [25]. However, cell lines differ from each other by many parameters. Here, we established a causal link between VHL inactivation, HIF1alpha stabilization and the increase of 2-hydroxyglutarate by performing a direct intervention.

Hypoxia and pseudohypoxia conditions, in which glycolysis is hyperactivated, present a challenge for a cell due to NADH accumulation. Interestingly, production of 2-hydroxyglutarate from 2-oxoglutarate by MDH consumes NADH. It can therefore be considered as one of the mechanisms to get rid of excess NADH that accumulates in glycolysis.

In the pioneering work of Thienpont et al. [7], the DNA hypermethylation was discovered in hypoxia. Under hypoxia, they observed a 50% increase in 2-hydroxyglutarate level and 5-10% increase of 2-hydroxyglutarate concentration normalized to the concentration of alpha-ketoglutarate, a cofactor of TET. However, this effect was much smaller than the direct effect of hypoxia and therefore DNA hypermethylation was attributed solely to the low concentration of oxygen required as a substrate for Tet enzymes. For comparison, in our study, 2HG/alpha-ketoglutarate ratio was 5-10 fold higher in Caki1*VHLmut* cells than in Caki1 cells (increased from 1 to 10%). Identical activity of Tet enzymes measured in nuclear extracts both in [7] between the cells in normoxia and hypoxia, and in our work (Figure S6B) between VHL wild-type cells and mutants can be explained by the fact that the assay used in both studies to estimate the rate of 5mC oxidation (Colorimetric Epigenase 5mC-Hydroxylase TET Activity/Inhibition Assay Kit from Epigentek) contained high molar excess of alpha-ketoglutarate that could neutralize the effect of 2HG increase. In other words, the kit attempts to measure Tet activity regardless of the concentrations of various metabolites in nuclear extracts.

The discovered link between activation of hypoxia signalling due to VHL inactivation, accumulation of 2-hydroxyglutarate and DNA hypermethylation not only explains hypermethylation in VHL mutants under normoxic condition, but can also contribute to DNA hypermethylation in hypoxia, additionally to the direct effect of low oxygen concentration on the activity of Tet enzymes. Deconvolution of these two effects requires further studies.

## Acknowledgements

This work was partially supported by Russian Scientific Fund grant №19-14-00347 for all authors except JS, MM, MG and DG who are members of University Center of Excellence “Towards Personalized Medicine” operating under Excellence Initiative – Research University in Nicolaus Copernicus University in Toruń.

## Data availability

Sequenced data was deposited to the GEO database (https://www.ncbi.nlm.nih.gov/geo/) under accession number GSE151787.

## Methods

### Cell culture

Caki-1 and HEK293t cell lines were grown in Dulbecco’s modified Eagle medium supplemented with 10% fetal bovine serum, 1% penicillin/streptomycin, and 2 mM L-glutamine. Cells were transfected with Lipofectamine 3000 (Thermo Fisher Scientific) according to the manufacturer’s recommendation.

### Genome editing

CRISPR/CAS9–based editing was performed as described in ref. [29]. Briefly, Caki1 cells were seeded on a 12-well plate. Cells were transfected with Lipofectamine 3000 with PX459-VHL plasmid to generate frame shift which led to a stop loss in the C-terminal alpha domain of VHL [3]. Puromycin was applied after 24 h at a concentration of 2 μg ml^−1^. Cells were incubated for 72 h, passaged into a 96-well plate at a density of 1 cell per 2 wells on medium without puromycin. For further study, we have chosen three clones for each experiment. The frame shift was confirmed by Sanger sequencing of the corresponding amplicons obtained from PCR with genomic DNA.

## Antibodies

Following antibodies were used in this study: anti-Hif1a ((H1 alpha 67) ab1), anti-actin (ab8227), anti-H3 (ab18521), anti-TET2 (active motif 61390).

### Viral vectors and lentivirus transduction

VHL-HA was subcloned to pCDF1-MCS2-EF1-Puro vector (System BioSciences). HEK293t cells were transfected with pCDF-VHL-HA, pVSV-G and pFiv-34N vectors. Viral supernatant was collected after 48 and 60 hrs, filtered through a 40um filter. Virus was precipitated by 12,5% PEG 8000 at +4^0^C overnight followed by centrifugation at 4200g during 40 minutes at+4^0^C. Pellet was resuspended in 300 mkl of OPTI-MEM and stored at -80^0^C. Infections were carried out in 4ml medium using polybrene diluted 1:2,000. 100ul of viral concentrate were used per 60-mm cell plate for 12 hrs. Infected cells were selected by puromycine (2 mkg/ml).

### Tumor xenografts

Nu/J mice were obtained from Jackson Laboratories and transferred to the Center for Genetic Resources of Laboratory Animals at the Institute of Cytology and Genetics, Siberian Branch, RAS (Stock No: 002019). All procedures were conducted under Russian legislation according to Good Laboratory Practice, inter-institutional bioethical committee guidelines and the European Convention for the protection of vertebrate animals used for experimental and other scientific purposes. All procedures were approved by the bioethical committee (protocol №33 from the 21.05.2018). All efforts were made to minimize the number of animals used and their suffering. All animals used had specific pathogen-free status. 5-6 mice were used per one cell line. Each mouse was inoculated subcutaneously in the withers area with 5 x 10^6^ Caki1, Caki1mutVHL or CakimutVHLexoVHL tumor cells in 100 mkl sterile PBS. Mice were sacrificed on day 21 after inoculation. Tumors were weighed and measured.

### Bisulfite conversion and whole-genome bisulfite sequencing

DNA methylation was profiled by whole genome bisulfite sequencing (WGBS). Two micrograms of genomic DNA from every cell line and 4 ng of lambda phage DNA were sonicated to average size 250 bp with Covaris S2. The end repair and an adenine was added to the 3′ end of the DNA fragments according to NebNext DNA UltraII protocol (NEB, USA). Preannealed forked Illumina adaptors containing 5′-methylcytosine instead of cytosine were ligated to both ends of DNA fragments using standard NEB adaptor ligation protocol (NEB, USA). Ligated fragments were treated with the sodium bisulfite and subsequent clean-up was performed with the EZ DNA Methylation™ Kit (ZymoResearch, USA) according to the manufacturer’s instructions. The bisulfite-treated DNA fragments were amplified using PCR and the following reaction: 5 µl of eluted DNA, 1 µl of NEB PE PCR two primers (1.0 and 2.0), and EpiMark® Hot Start Taq DNA Polymerase (NEB, USA). The amplification conditions were as follows: 5 min at 95 °C, 30 s at 98 °C then 15 (10 s at 98 °C, 30 s at 65 °C, 30 s at 72 °C), followed by 5 min at 72 °C. The PCR reaction was purified with AMPure XP beads (Beckman Coulter, USA), and final reduced representation bisulfite library was eluted in 30 µl EB buffer. The concentration of the final library was measured using the Agilent 2100 Bioanalyzer (Agilent Technologies, USA). The library was sequenced on Illumina 2500 platform according to standard Illumina cluster generation and sequencing protocols.

Reads were mapped to hg19 reference genome with Bismark software [30]. To estimate the efficacy of C-to-T conversion, lambda phage genome was added to the reference genome. Low DNA methylation levels at cytosines in lambda phage DNA indicate good bisulfite conversion. Additionally, we estimated DNA methylation level at C positions outside the CpG context. Differential methylation analysis was performed by MethPipe/RADmeth software package [31]. Custom scripts were used to iterate through bisulfite bam files and estimate global DNA methylation, per-chromosome methylation levels. To find out how DNA methylation changed in individual CpG islands, we calculated the total number of methylated and unmethylated CpG’s in bisulfite reads at each CpG position within the CpG island. Beta-binomial test was applied to estimate if the DNA methylation of a given CpG island significantly differed between two groups of samples.

The following genomic tracks were downloaded from ENCODE: histone marks for HepG2 cell line (wgEncodeBroadHistone tables, hg19 genome); clustered transcription factor binding sites, conserved across many cell lines (wgEncodeRegTfbsClusteredV3). PWM for HIF1a binding motif was downloaded from HOCOMOCO database [32,33]. Occurrences of HIF1a binding motif were searched in hg19 genome with SARUS software [34].

### RNA-seq

Total RNA was extracted from cell culture with Trizol reagent according to manufacture instruction. Quality was checked with BioAnalyser and RNA 6000 Nano Kit (Agilent). PolyA RNA was purified with Dynabeads® mRNA Purification Kit (Ambion). Illumina library was made from polyA RNA with NEBNext® Ultra™ II RNA Library Prep (NEB) according to manual. Sequencing was performed on HiSeq1500 with 50 bp read length. At least 10 millions of reads were generated for each sample.

Reads were mapped to hg19 genome with STAR software (version 2.5.4b), [35]. Gene models of non-overlapping exonic fragments were taken from the ENSEMBL 54 database. For each exonic fragment, total coverage by mapped reads in each sample was calculated with htseq-count (version 0.11.0), [36]. Differential expression analysis was performed by applying default read count normalization and performing per-gene negative binomial tests, implemented in DESeq2 R package (version 1.20.0), with default parameters [37].

### Flow cytometry

Cells were fixed in 70% ethanol during 30 minutes at +4^0^C, washed by 1xPBS 2 times, treated with ribonuclease and stained by PI and analyzed by flow cytometry. Data was analyzed by ModFit LT V5.09.

### TET activity assay

TET hydroxylase activity was measured by TET hydroxylase activity colorimetric quantification kit (ab15619) according to the manufacturer’s instruction.

### Mass-spectrometry profiling of modified nucleotides

The analyses were performed using a method described earlier by Gackowski et al. [38] with some modifications. Briefly, the pellet of the frozen cells was dispersed in ice-cold buffer B (Tris-HCl (10 mmol/L), Na_2_EDTA (5 mmol/L), deferoxamine mesylate (0.15 mmol/L), pH 8.0). SDS solution was added (to the final concentration of 0.5%), and the mixture was gently mixed using a polypropylene Pasteur pipet. Samples were incubated at 37 °C for 30 min. Proteinase K was added to the final concentration of 2.5 mg/mL and incubated at 37 °C for 1.5 h. The mixture was cooled to 4 °C, transferred to a centrifuge tube with phenol/chloroform/isoamyl alcohol (25:24:1), and vortexed vigorously. After extraction the aqueous phase was treated with a chloroform/isoamyl alcohol mixture (24:1). Supernatant was treated with two volumes of cold absolute ethanol in order to precipitate high molecular weight nucleic acids. The precipitate was removed with a plastic spatula and washed with 70% (v/v) ethanol and dissolved in MilliQ-grade deionized water. Samples were mixed with 200 mM ammonium acetate containing 0.2 mM ZnCl_2_, pH 4.6 (1:1 v/v). Nuclease P1 (1U) and tetrahydrouridine (10 μg/sample) were added to the mixture and incubated at 37°C for 1 h. Subsequently, 13 µL and 15 µL of 10% (v/v) NH_4_OH (for remaining nucleic acids and ultrafiltrate respectively) and 1.3 U of alkaline phosphatase were added to each sample following 1h incubation at 37 °C. Finally, all hydrolysates were ultrafiltered prior to injection. DNA hydrolysates were spiked with a mixture of internal standards in volumetric ratio 4:1, to concentration of 50 fmol/µL of [D_3_]-5-hmdC, [^13^C_10_, ^15^N_2_]-5-fdC, [^13^C_10_, ^15^N_2_]-5-cadC, and [13C10, 15N2]-5-hmdU. Chromatographic separation was performed with a Waters Acquity 2D-UPLC system with photo-diode array detector for the first dimension chromatography (used for quantification of unmodified deoxynucleosides and 5-mdC) and Xevo TQ-S tandem quadrupole mass spectrometer (used for second dimension chromatography and compounds analyzed in positive mode after first dimension: 5-hmdC and 8-oxodG, to assure better ionization at higher acetic acid concentration). At-column-dilution technique was used between the first and second dimension for improving retention at the trap/transfer column. The columns used were: a Waters Cortecs T3 column (150 mm×3 mm, 1.6 µm) with precolumn at the first dimension, a Waters X-select C18 CSH (100 mm×2.1 mm, 1.7 µm) at the second dimension and Waters X-select C18 CSH (20 mm×3 mm, 3.5 µm) as trap/transfer column. Chromatographic system operated in heart-cutting mode, indicating that selected parts of effluent from the first dimension were directed to trap/transfer column *via* 6-port valve switching, which served as “injector” for the second dimension chromatography system. The flow rate at the first dimension was 0.5 mL/min and the injection volume was 2 µL. The separation was performed with a gradient elution for 10 minutes using a mobile phase 0.05% acetate (A) and acetonitrile (B) (0.7-5% B for 5 minutes, column washing with 30% acetonitrile and re-equilibration with 99% A for 3.6 minutes). Flow rate at the second dimension was 0.3 mL/min The separation was performed with a gradient elution for 10 minutes using a mobile phase 0.01% acetate (A) and methanol (B) (1-50% B for 4 minutes, isocratic flow of 50% B for 1.5 minutes, and re-equilibration with 99% A up to next injection). All samples were analyzed in three to five technical replicates of which technical mean was used for further calculation. Mass spectrometric detection was performed using the Waters Xevo TQ-S tandem quadrupole mass spectrometer, equipped with an electrospray ionization source. Collision-induced dissociation was obtained using argon 6.0 at 3 x 10^−6^ bar pressure as the collision gas. Transition patterns for all the analyzed compounds, as well as specific detector settings were determined using the MassLynx 4.1 Intelli-Start feature in quantitative mode to assure best signal-to noise ratio and resolution of 1 at MS1 and 0.75 at MS2 (Supplementary Table 1).

### Mass-spectrometry profiling of metabolites

#### Quantitation of TCA metabolites: α-ketoglutaric acid (αKG), succinic acid (SA) and fumaric acid (FA)

Frozen cell culture samples (approximately 2-5×10^6^ cells) were suspended in 120 µL of double-distilled deionized water (Merck Millipore, Darmstadt, Germany) and sonicated for 5 min. After homogenization 20 µL of suspended cells were usued for further analysys of thymine. The rest of homogenate were used for measuring the level of TCA cycle metabolites (αKG, SA, FA) and were incubated in water bath at 100°C for 5 min. Then, 5 μl of 800 μmol/l isotope-labeled internal standards of [D_4_]-2-ketoglutaric acid (Sigma-Aldrich), [^13^C_4_]-succinic acid (Sigma-Aldrich) and [^13^C_4_]-fumaric acid (Sigma-Aldrich) were added to each sample. Subsequently, samples were purified by addition of 240 μl of waterless ethanol, vortexed and centrifuged at 24 400×*g* for 15 min at 4 °C. The supernatants (300 μl) were dried under vacuum for 2 hours and reconstituted in 100 μl deionized water (Merck Millipore, Darmstadt, Germany). The samples were filtered using AcroPrep Advance 96-Well Filter Plates 10 K and injected into ultra performance liquid chromatography system with tandem mass spectrometry detection (UPLC/MS/MS).

Waters Acquity UPLC consisted of binary gradient pump built-in vacuum degasser sample manager, column heater and PDA detector. The UPLC was operated using MassLynx 4.1 Software from Waters. Subsequent mass spectrometric detection was performed using a Waters Quattro Premier XE tandem quadrupole mass spectrometer, equipped with an electrospray ionization source. Chromatographic separation of examined metabolites (αKG, SA, FA) was achieved at 25°C using Synergy Hydro-RP C18 column (4 µm 80 Å 150 x 2 mm, Phenomenex) with a flow rate 0.25 mL/min and 3-µL injection volume. Formic acid (0.1%) and methanol were used as solvent A and B, respectively. The following program was used for αKG, SA and FA elution: 0–0.01 min. 99.9 % A, 0.1% B; 0.01–2.50 min. 92.0% A, 8% B; 2.50–3.50 min. 50% A, 50% B; 3.50–3.51 min. – 50% A, 50% B; 3.51–4.0 min – 99% A, 0.10% B. Multiple reaction monitoring (MRM) parameters were presented in Supplementary Table 2. The level of *TCA* cycle metabolites (αKG, SA, FA) in cell culture was determined as the ratio of the peak area derived from the isotopically labeled standard to the area under the peak from determining compound. Intracellular concentration of TCA metabolites (αKG, SA, FA) was calculated based on thymine level. All samples were analyzed at least in three technical replicates.

A series of standard curves were prepared over a concentration range of 0,391-100 µM from a stock solutions of unlabeled metabolites αKG (Sigma Aldrich), SA (Sigma Aldrich) and FA (Sigma Aldrich) in water and concentration range of 800 µM from a stock solutions of labeled metabolites [D_4_]-2-KG (Sigma-Aldrich), [^13^C_4_]-SA (Sigma-Aldrich) and [^13^C_4_]-FA (Sigma Aldrich) in water. All calibration curves constructed from the peak area ratios versus the molar ratios (labeled/unlabeled) showed a range of determination coefficient (R^2^) values over 0.99 (Supplementary Figure S10). The experiment was performed in five technical replicates.

#### Quantitation of L- and D-2-hydroxyglutarate

Frozen cell culture samples (approximately 2-5×10^6^ cells) were suspended in 200 µL of double-distilled deionized water (Merck Millipore, Darmstadt, Germany) and sonicated for 5 min. To 50 µL homogenate was added 2.5 µL mixture of 50 μM L-, D-[D_4_]-2-hydroxyglutaric acid (L-,D-[D_4_]-2HG) as internal standard (IS), 2.5 µL ammonium sulfate (10% v/v, POCH), 300 µL methanol (−80°C, LiChrosov) and vortexed. The stable-isotope labeled internal standards of L-, D-2-hydroxyglutaric acid-d6 were synthetized in-house by reduction of commercially available [D_6_]-2-ketoglutaric acid (Sigma-Aldrich) with excess zinc as previously described [39]. Samples were incubated at -80°C for 15 min. and centrifuged at 24 400×*g* for 15 min at 4 °C. The supernatants (350 μl) were dried under speedvac overnight. The dry residue was treated with 200 μl of freshly made 50 mg/ml (+)-Di-O-acetyl-L-tartaric anhydride (DATAN) in dichloromethane (Alfa Aescar) : acetic acid (Fischer Chemical) (4:1 by volume). The vial was capped and heated at 75 °C for 30 min. After the vial was cooled to room temperature, the mixture was evaporated to dryness under stream of nitrogen with XcelVap for 2 hours, redissolved in 30 μl deionized water (Merck Millipore, Darmstadt, Germany), filtrated using AcroPrep Advance 96-Well Filter Plates 10 K and injected into ultra-performance liquid chromatography system with Waters Quattro Premier XE tandem quadrupole mass spectrometer, equipped with an electrospray ionization source. The system was operated using MassLynx 4.1 Software from Waters. Chromatographic separation of examined L-, D-2HG enantiomers derivatives was achieved at 43°C using Acquity UPLC CSH C18 column (1.7 µm x 100 mm, Phenomenex) with a flow rate 0.35 mL/min and 0.5-µL injection volume. Ammonium formate (10 mM, pH 3.812, Fischer Chemical) and methanol were used as solvent A and B, respectively. The following program was used for L-,D-enantiomers derivatives elution: 0–0.10 min. 98.0 % A, 2.0% B; 0.10–2.00 min. 90.0% A, 10% B; 2.00–2.10 min. 70% A, 30% B; 2.10–3.30 min. – 70% A, 30% B; 3.30–3.31 min. 98% A, 2.00-% B; 3.31-5.00 min. 98% A, 2.00% B. Product ion transitions was monitored at 363.2>147.1 and 367.2>151.1 for examined L-, D-2-hydroxyglutarate and stable-isotope-labeled L-,D-[D_4_]-2HG derivatives (IS), respectively.

The level of L-, D-2HG enantiomers in cell culture was determined as the ratio of the peak area derived from the isotopically labeled standard to the area under the peak from determining compound. The results were normalized “per cell” on the basis of concentration of thymine in the homogenate. Thymine level was measured using UPLC-UV technique after acidic hydrolysis of the sample. All samples were analyzed at least in three technical replicates. A series of standard curves were prepared over a concentration range of 0.005-10 µM from a stock solutions of unlabeled enantiomers L-, D-2HG (Sigma Aldrich) in water and 50 µM of labeled metabolites L-,D-[D_4_]-2HG. All calibration curves constructed from the peak area ratios versus the molar ratios (labeled/unlabeled) showed determination coefficient (R^2^) values over 0.99 (Supplementary Figure S11).

