## Supplementary figures for "An IDH-independent mechanism of DNA hypermethylation upon VHL inactivation in cancer"

2-hydroxyglutarate drives whole-genome hypermethylation in kidney cancer cells with inactivated VHL

### Supporting Information

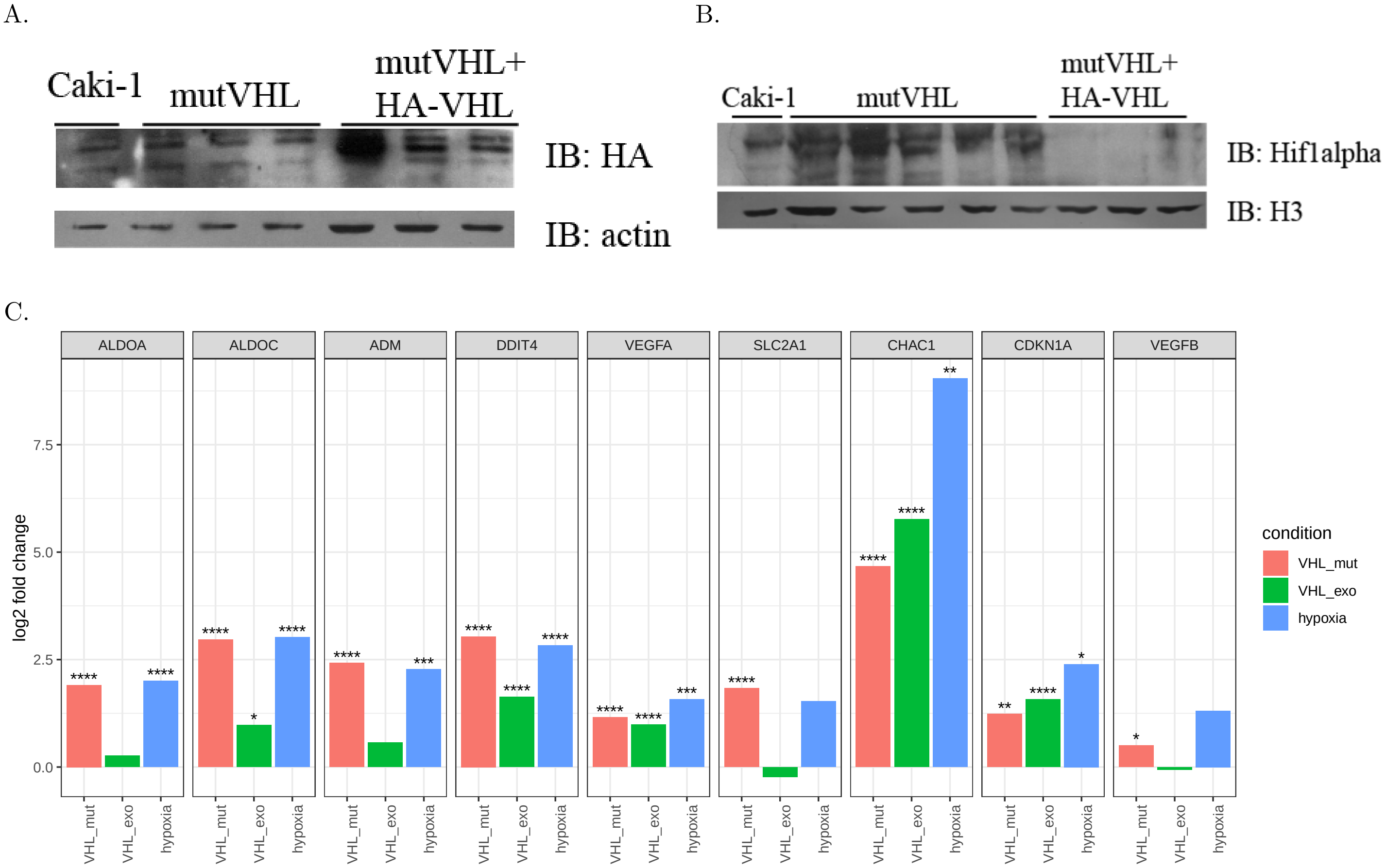


**Figure S1.** (A) Western blot for HA tag that marks expression of exogenous VHL gene and (B) Hif1$\alpha$ protein. Inactivation of VHL induced accumulation of Hif1$\alpha$ protein. The level of Hif1$\alpha$ protein decreased after exogenous re-activation of VHL. (C). Changes of gene expressions of genes known to be activated under hypoxia. VHL inactivation caused changes similar to hypoxia. These changes were parially reverted by exogenous expression of wild-type VHL. Y-axis indicates log fold-change of gene expression in VHL mutants, rescued VHL mutants with exogenic expression of VHL and Caki-1 cells in hypoxic conditions, all compared to Caki-1 cells in normoxic conditions. DESeq2 p-values corrected for multiple testing (FDR) are indicated as follows: **** $P < 0.0001$; *** $P < 0.001$; ** $P < 0.01$; * $P < 0.05$.

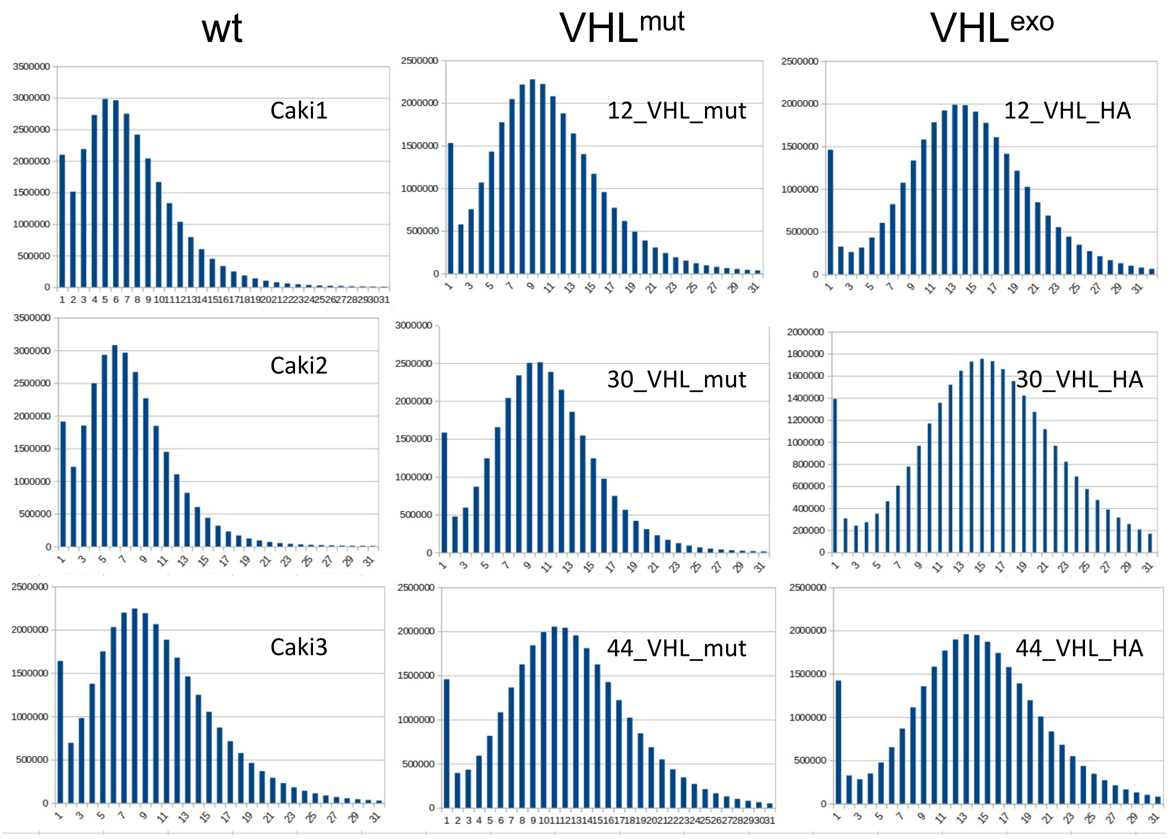


**Figure S2.** Distributions of genomic coverage by bisulfite reads for individual CpG positions in all studied samples.

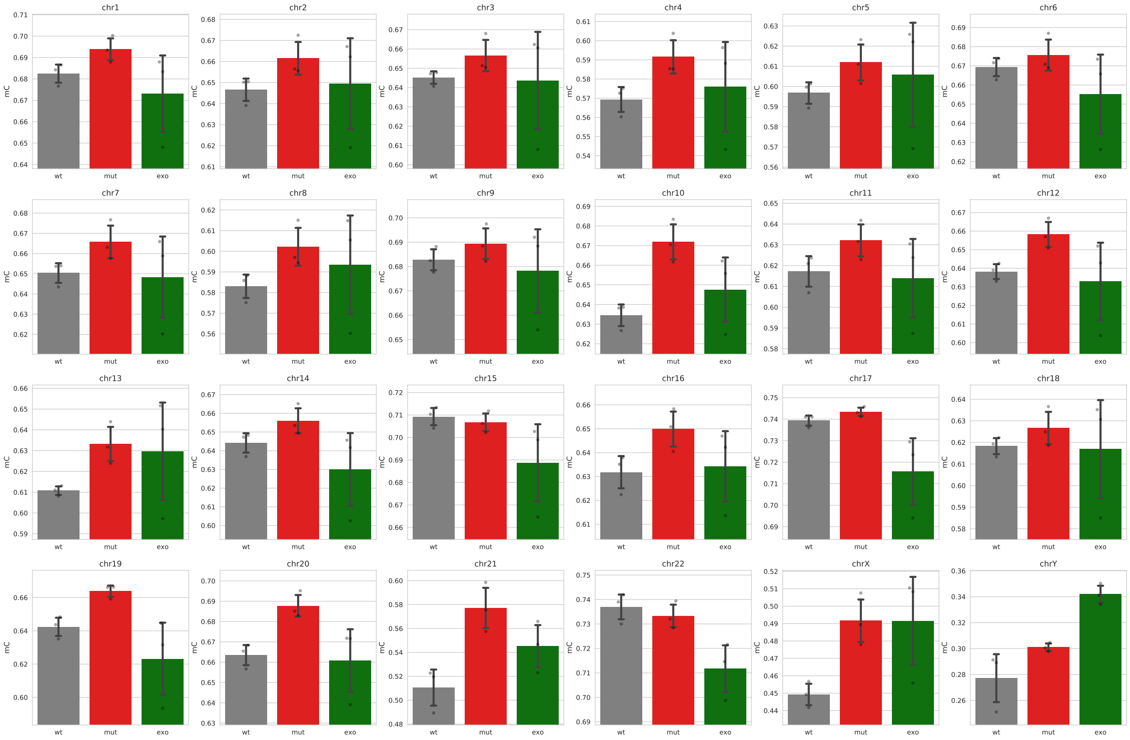


**Figure S3.** Average DNA methylation level per chromosome in Caki-1 cells (wt), VHL mutants (mut) and VHL mutants with rescued VHL expression (exo)

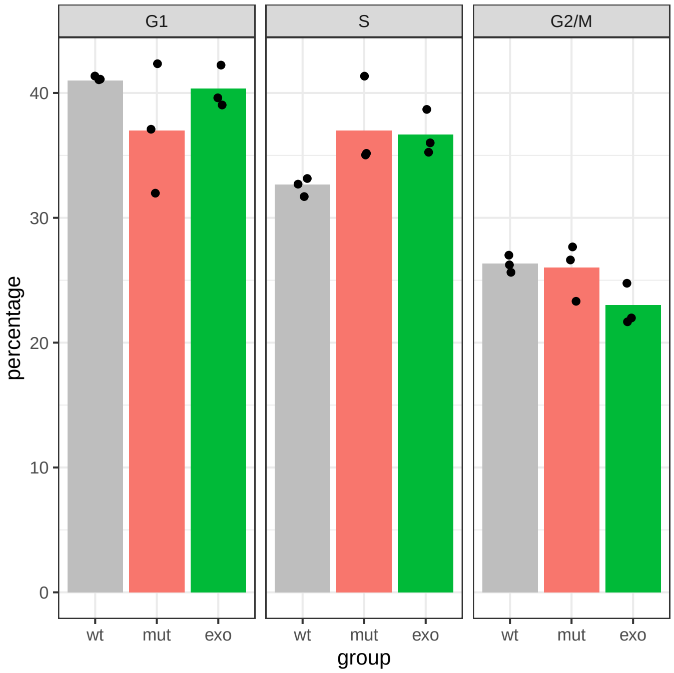


**Figure S4.** Distribution of the cells according to the phases of the cell cycle. No significant changes were observed between wild-type and VHL mutant clones: according to Wilcoxon–Mann–Whitney test, $P = 0.64$ for the comparison of the fractions of cells in the G1 phase and $P = 0.07$ for the fractions of cells in the S phase.

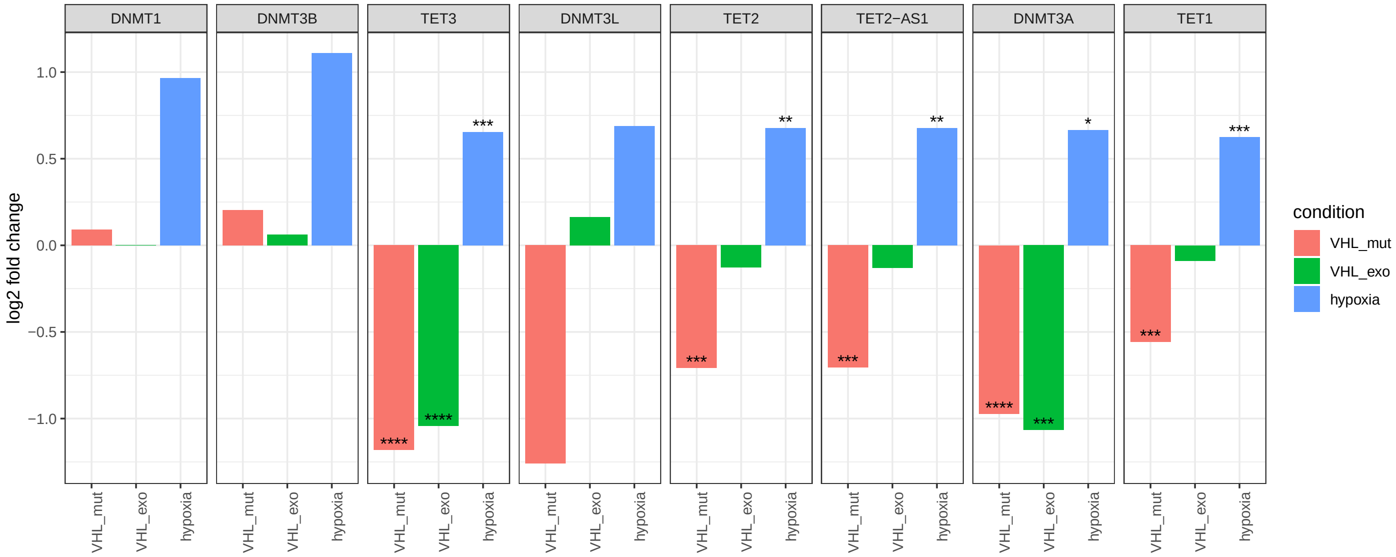


**Figure S5.** Changes of gene expressions of TET genes and *de novo* DNA methyltransferases (DNMTs). Y-axis indicates log fold-change of gene expression in VHL mutants, rescued VHL mutants with exogenic expression of VHL and Caki-1 cells in hypoxic conditions, all compared to Caki-1 cells in normoxic conditions. DESeq2 p-values corrected for multiple testing (FDR) are indicated as follows: **** $P < 0.0001$; *** $P < 0.001$; ** $P < 0.01$; * $P < 0.05$.

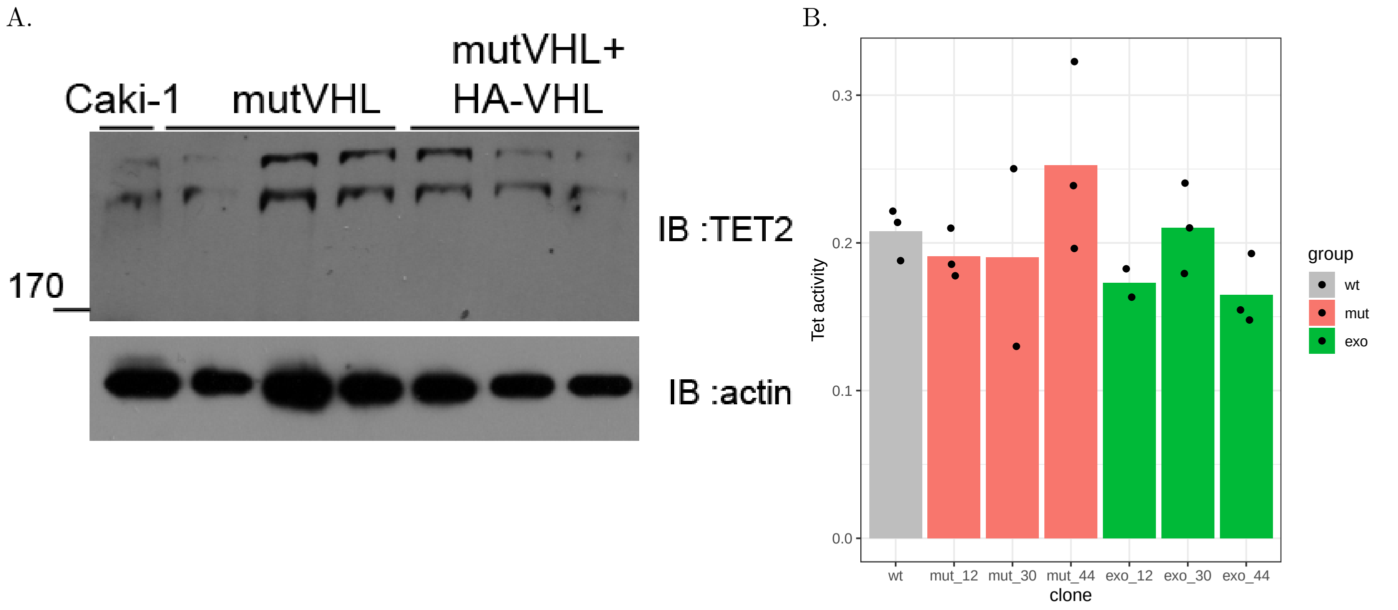


**Figure S6.** (A) Western blot for Tet2 protein. (B) Calorimetric analysis of Tet activity

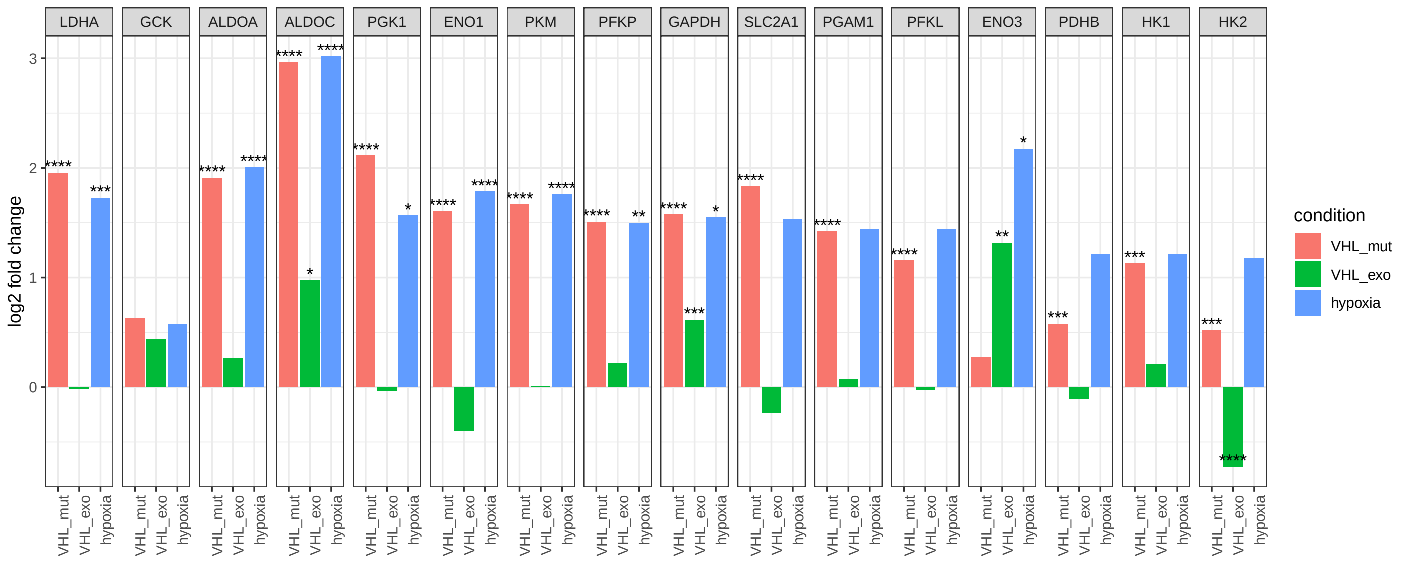


**Figure S7.** Changes of gene expressions of glycolysis genes. Y-axis indicates log fold-change of gene expression in VHL mutants, rescued VHL mutants with exogenic expression of VHL and Caki-1 cells in hypoxic conditions, all compared to Caki-1 cells in normoxic conditions. DESeq2 p-values corrected for multiple testing (FDR) are indicated as follows: **** $P < 0.0001$; *** $P < 0.001$; ** $P < 0.01$; * $P < 0.05$.

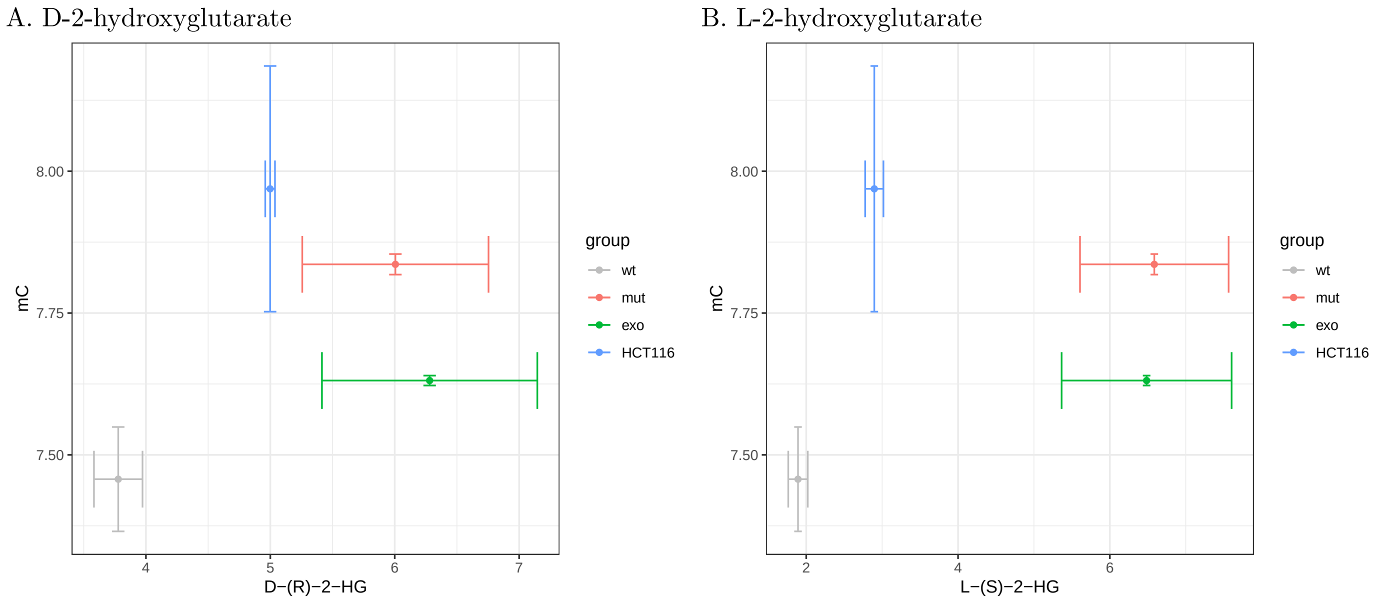


**Figure S8.** Concentrations of (A) D-2-hydroxyglutarate and (B) L-2-hydroxyglutarate (x-axis) and average DNA methylation level (y-axis) in Caki-1 cells (wt), VHL mutants (mut), VHL mutants with rescued VHL expression (exo) and HCT116 cell line with known IDH gain-of-function mutation

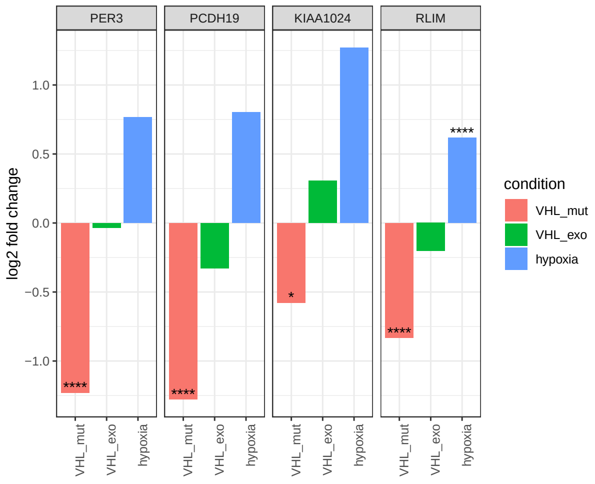


**Figure S9.** Changes of gene expressions of tumor suppressor genes with promoters that were hypermethylated in VHL mutants. Y-axis indicates log fold-change of gene expression in VHL mutants, rescued VHL mutants with exogenic expression of VHL and Caki-1 cells in hypoxic conditions, all compared to Caki-1 cells in normoxic conditions. DESeq2 p-values corrected for multiple testing (FDR) are indicated as follows: **** $P < 0.0001$; *** $P < 0.001$; ** $P < 0.01$; * $P < 0.05$.

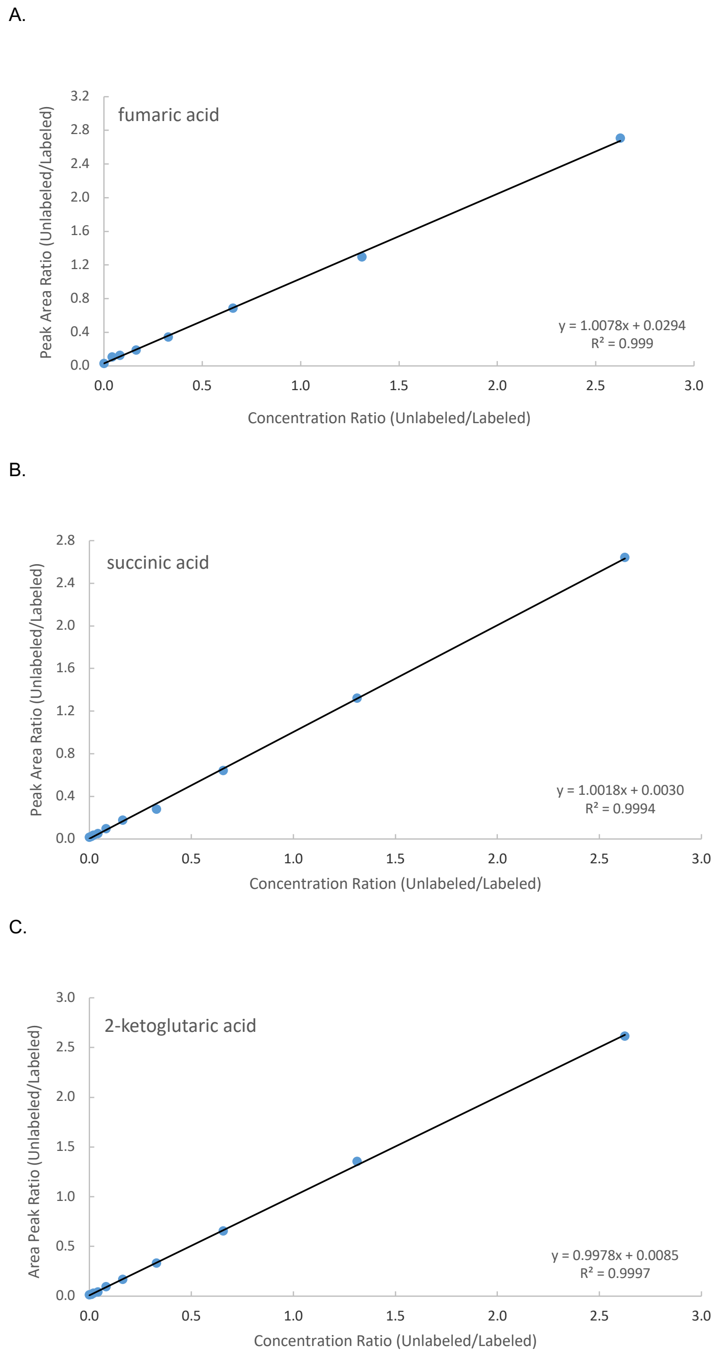


**Figure S10.** (A). Calibration curve of fumaric acid, obtained from the measured ratio values of the peak area of unlabeled FA to the peak area of labeled $[^{13}\mathbf{C}_4]\mathbf{-FA}$ vs. ratio of the unlabeled FA to labeled $[^{13}\mathbf{C}_4]\mathbf{-FA}$ concentrations. (B). Calibration curve of succinic acid, obtained from the measured ratio values of the peak area of unlabeled SA to the peak area of labeled $[^{13}\mathbf{C}_4]\mathbf{-SA}$ vs. ratio of the unlabeled SA to labeled $[^{13}\mathbf{C}_4]\mathbf{-FA}$ concentrations. (C). Calibration curve of 2-ketoglutaric acid, obtained from the measured ratio values of the peak area of unlabeled KG to the peak area of labeled $[\mathbf{D}_4]-\mathbf{KG}$ vs. ratio of the unlabeled 2-KG to labeled $[\mathbf{D}_4]-\mathbf{KG}$ concentrations.

A.

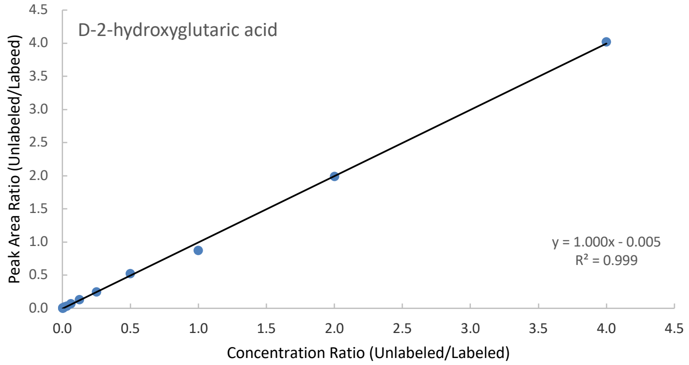


B.

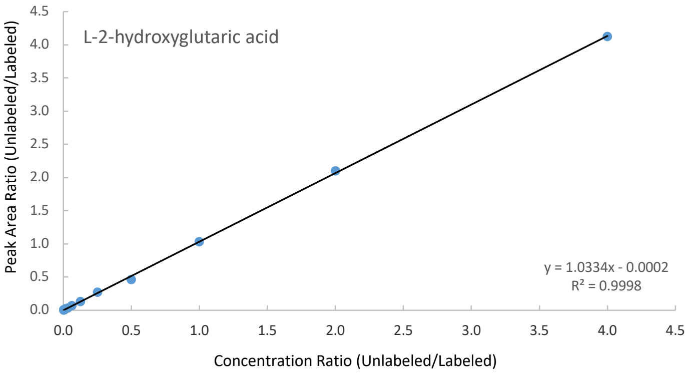


**Figure S11.** (A). Calibration curve of D-2-hydroksyglutaric acid, obtained from the measured ratio values of the peak area of unlabeled D-2-HG to the peak area of labeled $D-[\mathbf{D}_4]-\mathbf{2}-\mathbf{HG}$ vs. ratio of the unlabeled D-2-HG to labeled $D-[\mathbf{D}_4]-\mathbf{2}-\mathbf{HG}$ concentrations. (B). Calibration curve of L-2-hydroksyglutaric acid, obtained from the measured ratio values of the peak area of unlabeled L-2-HG to the peak area of labeled $L-[\mathbf{D}_4]-\mathbf{2}-\mathbf{HG}$ vs. ratio of the unlabeled L-2-HG to labeled $D-[\mathbf{D}_4]-\mathbf{2}-\mathbf{HG}$ concentrations.
